# Spinal microglia are dispensable for the development of opioid-induced hyperalgesia

**DOI:** 10.64898/2026.09.24.753592

**Authors:** Julia E. R Nickols, Tom Wilson, Kelly V. Lee, Jayme M. Clark, Adam C. Wass, Charbel S. Baaklini, Christian Faig, Madelene F.S. Ho, Katherine M. Souter, Bozena Szulc, Marianela E. Traetta, Jason R. Plemel, Anna M.W. Taylor

**Author notes:** **Correspondence**: Anna M.W. Taylor, Ph.D., Department of Pharmacology, University of Alberta, Edmonton, Alberta, Canada T6G 2R3.

## Abstract

Adverse side effects of chronic opioid use, including analgesic tolerance, opioid-induced hyperalgesia, and addiction, are serious clinical challenges for pain management and opioid deprescribing. Microglia have been implicated in the development of these adverse effects, but the extent to which opioids alter microglia and the mechanisms behind opioid effects on microglia are poorly understood. We performed morphological and single-cell transcriptomic analysis on spinal microglia from opioid-withdrawn and saline-treated mice, and found that while morphological changes recapitulated prior reports, transcriptomic changes in these cells were modest. Furthermore, peripherally, but not centrally, restricted opioid administration was sufficient to induce both hyperalgesia and microglial morphological changes in mice. Finally, complete loss of microglia in *Csf1r*^Δ/ΔFIRE^ mice did not prevent the development of opioid-induced hyperalgesia, analgesic tolerance, and somatic withdrawal symptoms. These data argue against any microglial involvement in the development of these phenomena.

## Introduction

Opioids are well-known and widely used analgesics. However, chronic opioid use results in a paradoxical pain hypersensitivity, known as opioid-induced hyperalgesia (OIH) (1–3). OIH is particularly problematic in both chronic pain (4,5) and substance use disorder populations (6–8). In the latter, pain perception in withdrawal is associated with increased drug craving (9), riskier injection practices (10), and increased risk of opioid overdose (11).

The notion that opioids produce nociceptive hypersensitivity has long been recognized in animal models (12). The mu opioid receptor (MOR) is widely distributed throughout both the central and peripheral nervous systems (13–17), and OIH, alongside other withdrawal-associated symptoms, is MOR-dependent (18,19). However, the exact MOR-expressing cellular populations that are required to produce and maintain OIH remain unclear. MOR expression in primary afferents (20–23), spinal interneurons (24), and supraspinal circuits (25,26) have all been implicated in the development of OIH.

Others, however, have suggested a role for glial reactivity in the emergence and development of OIH, with a particular focus on spinal microglia and astrocytes. Increased glial reactivity and morphological changes are noted to co-occur with chronic opioid administration and withdrawal (27), and the idea that microglia in particular may contribute to the development of OIH has been an attractive theory, given the noted roles of these cells in pain modulation via the release of inflammatory mediators, neurotrophic factors, and the pruning of neuronal synapses and perineuronal nets (28). In animal models, microglia have been implicated in the development of both analgesic tolerance and OIH (29–35), as well as somatic withdrawal symptoms (36). Additionally, the activation of toll-like receptor 4 (TLR4) by opioid agonists has received considerable attention as a potential route through which opioid-induced microglial reactivity may occur (37–40). In line with this, single-nucleus sequencing of the brains of opioid-use disorder patients has largely described enhanced pro-inflammatory signaling in microglia (41,42).

However, conflicting evidence exists about both the extent to which microglia are altered following chronic opioid exposure and the functional outcomes of these alterations: both TLR4 (43,44) and MOR (45) expression in microglia appears to be largely dispensable for both OIH and analgesic tolerance, and the glial inhibitor minocycline fails to improve pain in patients maintained on opioid agonist treatment (46). Furthermore, a more recent study (47) described no changes in the development or maintenance of OIH in mice when microglia were depleted with either a CSF1R inhibitor or genetic targeting of CD11b^+^ cells, despite earlier reports of OIH being abrogated by glial inhibitors (31,34,44) and antibody-mediated targeting of CD11b^+^ cells (30). Notably, depletion efficacy varied substantially between these studies, leaving open the possibility that residual microglia may still contribute to OIH.

Here, we demonstrate that while chronic opioid administration in mice produces persistent OIH and alterations in the morphology of spinal cord microglia, these changes are uncoupled to any evidence of overt microglial reactivity. Furthermore, single-cell RNA sequencing of spinal cord microglia in withdrawal demonstrated only a modest and transient increase in the reactivity of these cells in the acute withdrawal period. Direct exposure of microglia to opioids *in vivo* was not required for the development of OIH, and a complete loss of microglia did not prevent the development of OIH, analgesic tolerance, or somatic withdrawal symptoms in mice.

## Results

### Escalating morphine produces prolonged hyperalgesia associated with morphological changes in microglia

We first characterized the emergence and persistence of opioid-induced hyperalgesia following an escalating morphine dosing regimen (Fig. 1A). Acutely withdrawn mice (day 1 post-treatment) demonstrated both mechanical (Fig. 1B) and thermal (Fig. 1C) hypersensitivity, with mechanical, but not thermal, hypersensitivity resolving by 1-week post-drug. We confirmed the persistence of thermal hypersensitivity using the thermal gradient test (Fig. 1D), where mice were placed on rectangular platform with opposite ends set to either noxious hot (> 42 °C) or noxious cold (<17 °C). At both 1- and 8-days post-drug, opioid-withdrawn mice displayed heightened avoidance of the noxious temperature zones relative to saline controls. Given previous evidence of spinal microglial involvement in mediating opioid-induced hyperalgesia, we examined the state of spinal microglia during this 7-day post-drug period when mice demonstrate pain hypersensivity. Acute morphine withdrawal failed to significantly alter the IBA1^+^ area in the spinal dorsal horn (Fig. 1E), and neither acute (Fig. 1F) or prolonged (Fig. S1A, B) morphine withdrawal altered the proportion of spinal cord microglia/macrophages. Acute and prolonged morphine withdrawal failed to increase protein expression of CD45, CD11b, or IBA1 by microglia (Fig. 1G, Fig. S1C-E), and similarly the expression of CD11b mRNA (*Itgam*) was unchanged in the spinal cord at both day 1 and 7 post-drug (Fig. S1F). No upregulation of the classic proinflammatory cytokines *Il1b*, *Il6*, or *Tnf* was observed in spinal cords of mice at day 1 post-morphine (Fig. S1G). On the other hand, the morphology of dorsal horn microglia was significantly altered at day 1 post-morphine (Fig. 1H,I), with acutely withdrawn microglia demonstrating soma enlargement and reduced branching, the latter of which was maintained at day 7 post-drug (Fig. 1J). Overall, as previously described, we observed consistent morphological changes in spinal microglia, but these were uncoupled to other metrics of overt glial reactivity, including the expression of microglia-associated surface proteins and microgliosis.

**Figure 1.**
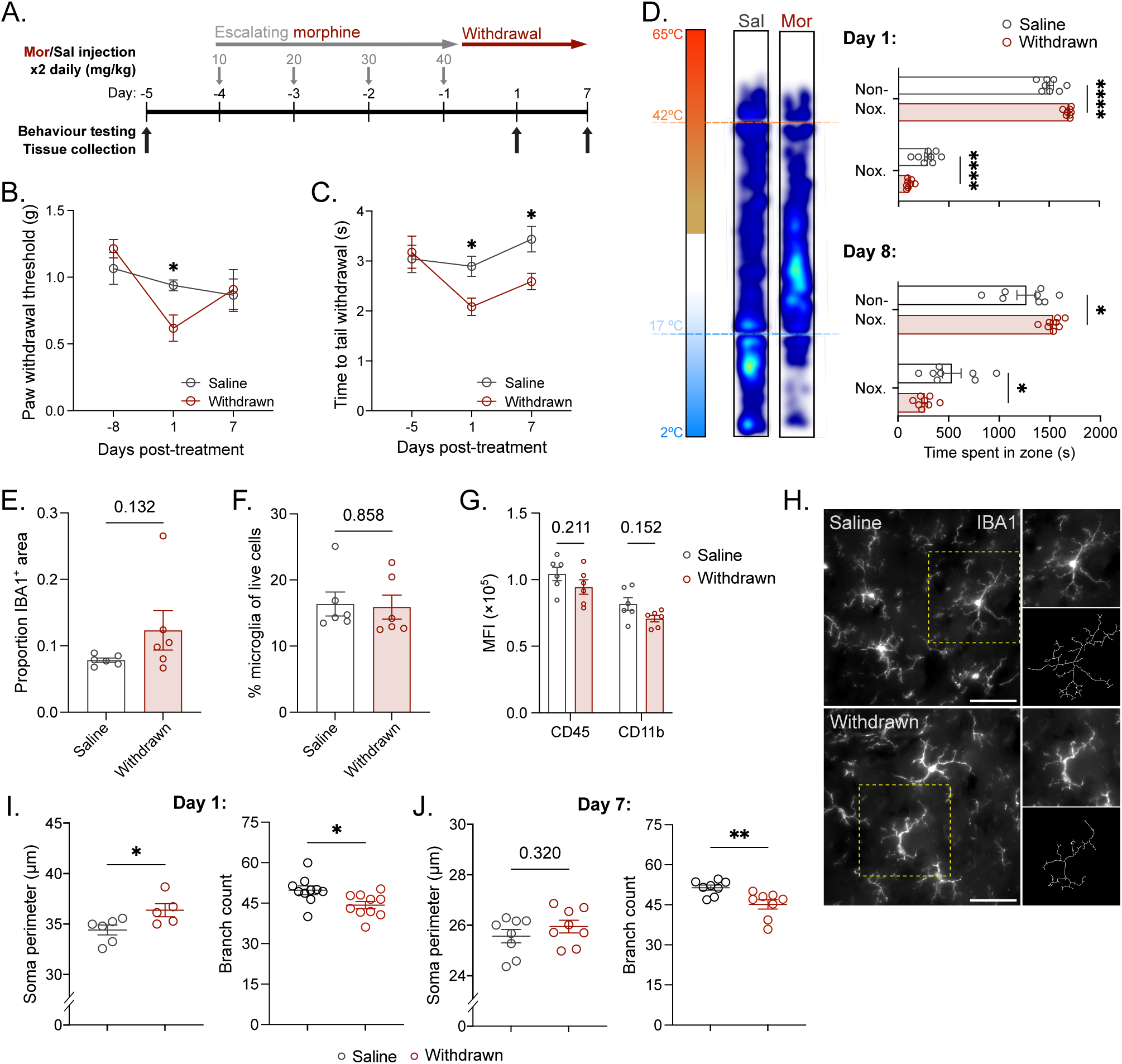
Chronic morphine exposure produces prolonged hyperalgesia and alterations in microglial morphology. (A) Opioid dosing regimen. (**B** - **D**) Chronic escalating opioid treatment produces mechanical hyperalgesia in a Von Frey assay (B) (n_M_ = 8 /group), and thermal hyperalgesia in both tail withdrawal (C) (n_M_ = 4, n_F_ = 4/group) and thermal gradient (D) (n_M_ = 8 /group) tests. (**E** - **G**) Chronic opioid treatment did not significantly increase total IBA1^+^ area in the spinal dorsal horn at day 1 post-morphine (E) (n_M_ = 3-4, n_F_ = 2/group) and does not alter the proportion of spinal cord microglia (CD45^+^CD11b^+^Gr1^-^) (F) or the median fluorescent intensity (MFI) of CD45 and CD11b in these cells (F) (n_M_ = 4, n_F_ = 2/group). (**H**) Representative images of dorsal horn spinal cord microglia in saline-treated and opioid-withdrawn mice. Scale bar = 50 µm. (**I** and **J**) Skeleton analysis of individual dorsal horn microglia in saline-treated and morphine-withdrawn mice revealed increases in soma size and decreases in branch counts at day 1 post-drug (I) (n_M_ = 3-8, n_F_ = 2/group) and decreases in branch counts at day 7 post-drug (J) (n_M_ = 8/group). Data is represented as mean ± SEM, two-way repeated measures ANOVA with Sidak-Holm post-hoc (B, C); two-way ANOVA with Sidak-Holm post-hoc (D); independent t test (I, J) with Welch correction (F); Mann-Whitney U test (E); multiple independent t tests with Sidak-Holm correction (G). *p<0.05, **p<0.01, ***p<0.001, ****p<0.0001.

### Spinal microglia demonstrate minimal transcriptomic changes during morphine withdrawal

To determine whether morphological changes in spinal microglia may be coupled to an altered transcriptional state, we isolated live nucleated CD11b^+^ cells from the lumbar spinal cords of saline-treated and morphine-withdrawn mice at days 1 and 7 post-drug for single-cell RNA sequencing (scRNA-seq) (Fig. 2A). Following quality control (Fig. S2) and the removal of non-microglial CD11b^+^ cells (*Tmem119*^-^*P2ry12*^-^clusters), we clustered the remaining microglia (15,809 cells) to define five distinct transcriptional states in spinal cord microglia (Fig. 2B, C). This included two homeostatic states (HM 1 and 2), defined by high expression of complement and ribosomal genes (HM 1; high *C1qb*, *Fcer1g*, *Rplp0* expression) and high expression of genes related to cell motility and cytoskeletal rearrangement (HM 2; high *Elmo1*, *Dock4*, *Pde3b* expression), respectively. We also identified clusters of cytokine-reactive microglia (CRM; high *Ccl4*, *Ccl3*, *Tnf* expression), interferon-response microglia (IRM; high *Cxcl10*, *Mx1*, *Isg15* expression), and microglia with expression profiles similar to previously described disease-associated microglia (48,49) (DAM; high *Apoe*, *Clec7a*, *Lyz2* expression). Notably, the CRM cluster also displayed relatively high expression of some DAM-related genes, including *Apoe* and *Lpl*, indicating some mixing of DAM-like and inflammatory microglia in this cluster. All identified clusters were present across all samples, regardless of treatment, suggesting that all five identified states represent the typical transcriptional heterogeneity of spinal cord microglia.

**Figure 2.**
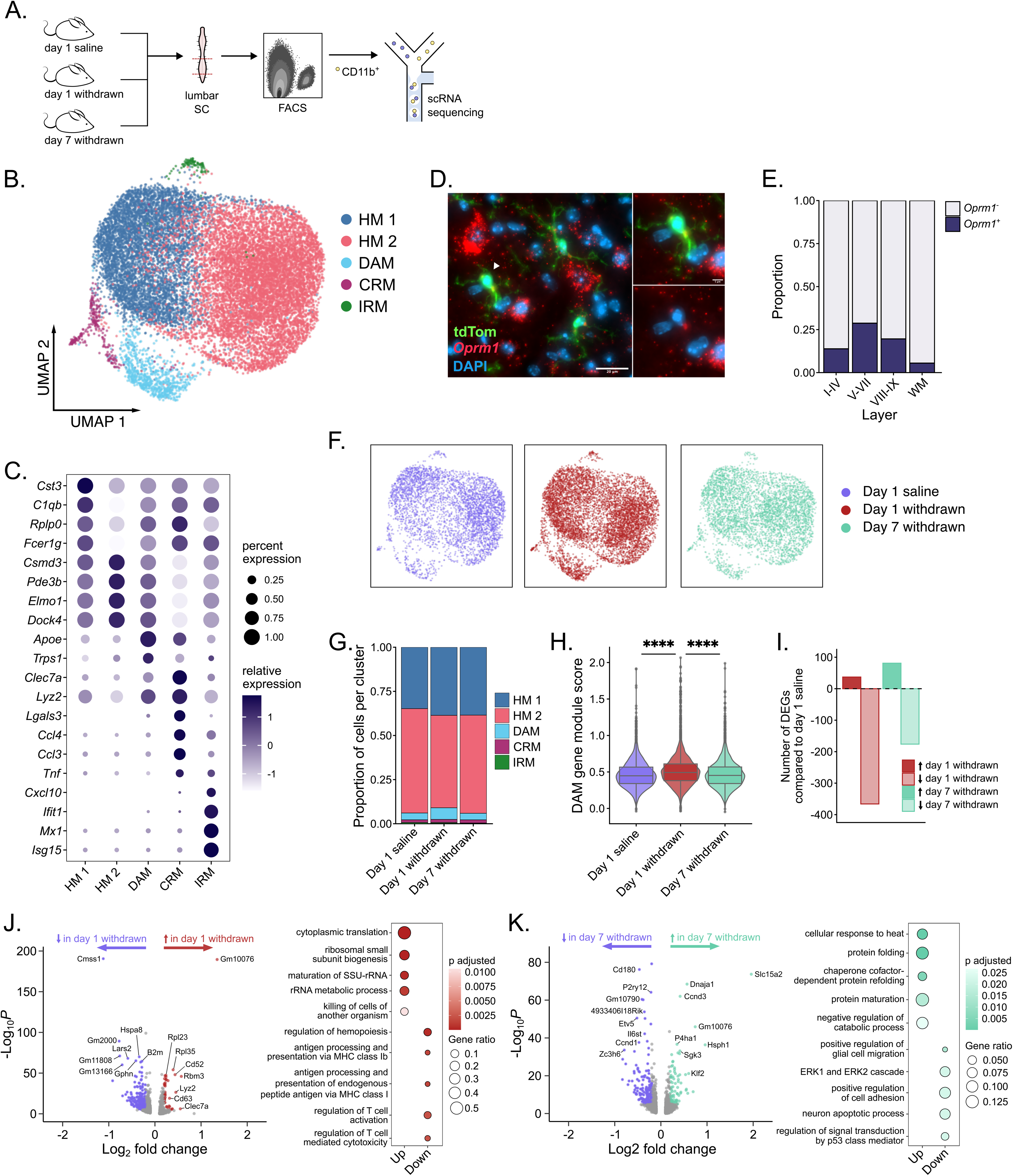
Single-cell RNA sequencing of lumbar spinal cord microglia in saline-treated and opioid-withdrawn mice. (**A**) Live nucleated CD11b^+^ spinal cord (SC) cells were sorted via fluorescence activated cell sorting (FACS) and sequenced on a 10x Genomics platform. (**B**) UMAP and clustering of all recovered microglia following quality control, with spinal cord microglia sorted into 2 homeostatic populations (HM 1 and 2), disease-associated microglia (DAM), cytokine-reactive microglia (CRM), and interferon-response microglia (IRM). (**C**) Top gene markers for each of the established microglial clusters. (**D**) In-situ hybridization for *Oprm1* in the spinal cords of naïve CX3CR1-tdTomato (tdTom) mice. (**E**) Proportions of *Oprm1*^+^ and *Oprm1*^-^microglia in spinal layers I-IX and in white matter (WM) regions. (**F**) UMAPs of microglia isolated from mice at 1-day post-saline or opioid treatment, or 7 days post-opioid treatment. (**G**) Proportions of microglia from each condition in each microglial cluster. (**H**) Gene module scores for canonical DAM genes in cells from each of the three treatments. (**I**) Number of up- and down-regulated genes in day 1 and day 7 withdrawn compared to day 1 saline. (**J** and **K**) Volcano plots (minimum log_2_ fold change > 0.2 and p_adjusted_ < 0.05) and biological processes gene ontology (GO) terms following over-representation analysis for differentially expressed genes in day 1 withdrawn (J) and day 7 withdrawn (K) relative to day 1 saline. ****p<0.0001.

Conflicting prior reports (20,50,51) exist regarding whether the mu opioid receptor is expressed in mouse microglia *in vivo*. In our data, we detected the presence of mRNA encoding the mouse mu opioid receptor (*Oprm1*; ∼28.9% of all microglial cells) within microglia in our scRNA-seq dataset. On the other hand, we failed to detect expression of the other canonical opioid receptors, including *Oprk1*, *Oprd1*, and *Oprl1* in spinal microglia. Similar to what has been previously described (20), we found that the majority of reads attributed to *Oprm1* mapped to intronic regions, with few reads in exons 1-3, which encode the transmembrane domains of the canonical MOR (Fig. S3A). To confirm the expression of *Oprm1* by microglia, we performed in situ hybridization for mature *Oprm1* mRNA in the spinal cords of drug-naïve *Cx3cr1*^CreER +/−^; ROSA26^tdT/-^mice (Fig. 2D, E). Overall, *Oprm1* expression in microglia was modest, but detectable in a subset of cells. The highest proportion of *Oprm1*^+^ CX3CR1^+^ microglia was detected in layers V-VIII of the spinal cord (∼28.7%), with the lowest proportion of *Oprm1*^+^ CX3CR1^+^ microglia detected in white matter regions (∼5.5%).

Despite some microglial *Oprm1* expression, opioid-withdrawn microglia at both 1- and 7-days post-drug displayed relatively modest transcriptional changes relative to day 1 saline controls (Fig. 2F-K). The distribution of microglia across clusters was largely unchanged in both day 1 and 7 withdrawn samples, apart from a small increase in the proportion of microglia residing in the DAM cluster in the day 1 withdrawn (6.8%) compared to the day 1 saline (3.8%) samples (Fig. 2G, S3B). To confirm that our DAM cluster was comparable to previously described DAM, we scored the cells in our dataset on a gene module comprised of conserved DAM markers across 12 microglial transcriptomics studies in humans and mice (52) and observed a significant increase in the DAM gene module score among day 1 withdrawn microglia (Fig. 2H). Despite the small increase in the proportion of DAM, overall opioid withdrawal resulted in a relatively small number of upregulated differentially expressed genes (DEGs) (Fig. 2I; 37 in day 1 withdrawn; 81 in day 7 withdrawn), with a larger proportion of DEGs downregulated relative to day 1 saline (366 in day 1 withdrawn; 176 in day 7 withdrawn). Similarly, pseudobulking of replicates resulted in only a small set of DEGs in day 1 withdrawn mice, and no differentially expressed genes at day 7 post-drug (Fig. S3C-D). In line with the increase in DAM in the day 1 withdrawn group, the DAM-associated genes *Lyz2* and *Cd63* were significantly upregulated in day 1 withdrawn samples in both the single-cell and pseudobulked DEG testing. The DAM state is highly associated with microglial phagocytosis, particularly of myelin debris (53–55). Given their expression of *Oprm1*, we questioned whether opioids may directly shift microglia towards a DAM-like state. Direct exposure of primary mouse microglia to morphine, however, failed to increase engulfment of either myelin or nuclear debris *in vitro* (Fig. S4A-D). Consequently, despite their expression of the *Oprm1* gene, spinal microglia are only minimally altered by acute and prolonged opioid exposure.

### Peripheral, but not central, opioid exposure produces hyperalgesia in mice

To determine whether central, and particularly microglial, opioid exposure was required to produce opioid-induced hyperalgesia in mice, we treated mice chronically with the mu opioid agonist DAMGO, a blood-brain barrier impermeable opioid peptide (Fig. 3A). DAMGO administered via the i.p. route at a dose of 6 mg/kg produced modest acute analgesia (Fig. 3B), while failing to elicit the locomotor effects of centrally acting mu agonists (Fig. S5A). Chronic peripheral DAMGO exposure elicited mechanical hypersensivity in mice upon withdrawal (Fig. 3C), demonstrating that direct microglial opioid exposure is dispensable for pain hypersensivity following opioid treatment. Conversely, repeated i.c.v. injections of DAMGO in mice (Fig. 3D) resulted in robust acute analgesia (Fig. 3E) but failed to alter mechanical thresholds during acute withdrawal (Fig. 3F). Interestingly, repeated i.p., but not i.c.v., injections of DAMGO also produced a modest but significant decrease in the number of projections observed in spinal dorsal horn microglia (Fig. 3G-I, Fig. S5B-I), suggesting that direct opioid exposure on microglia is not only dispensable for opioid-induced hyperalgesia, but also may be dispensable for some of the morphological alterations observed following systemic morphine exposure and withdrawal.

**Figure 3.**
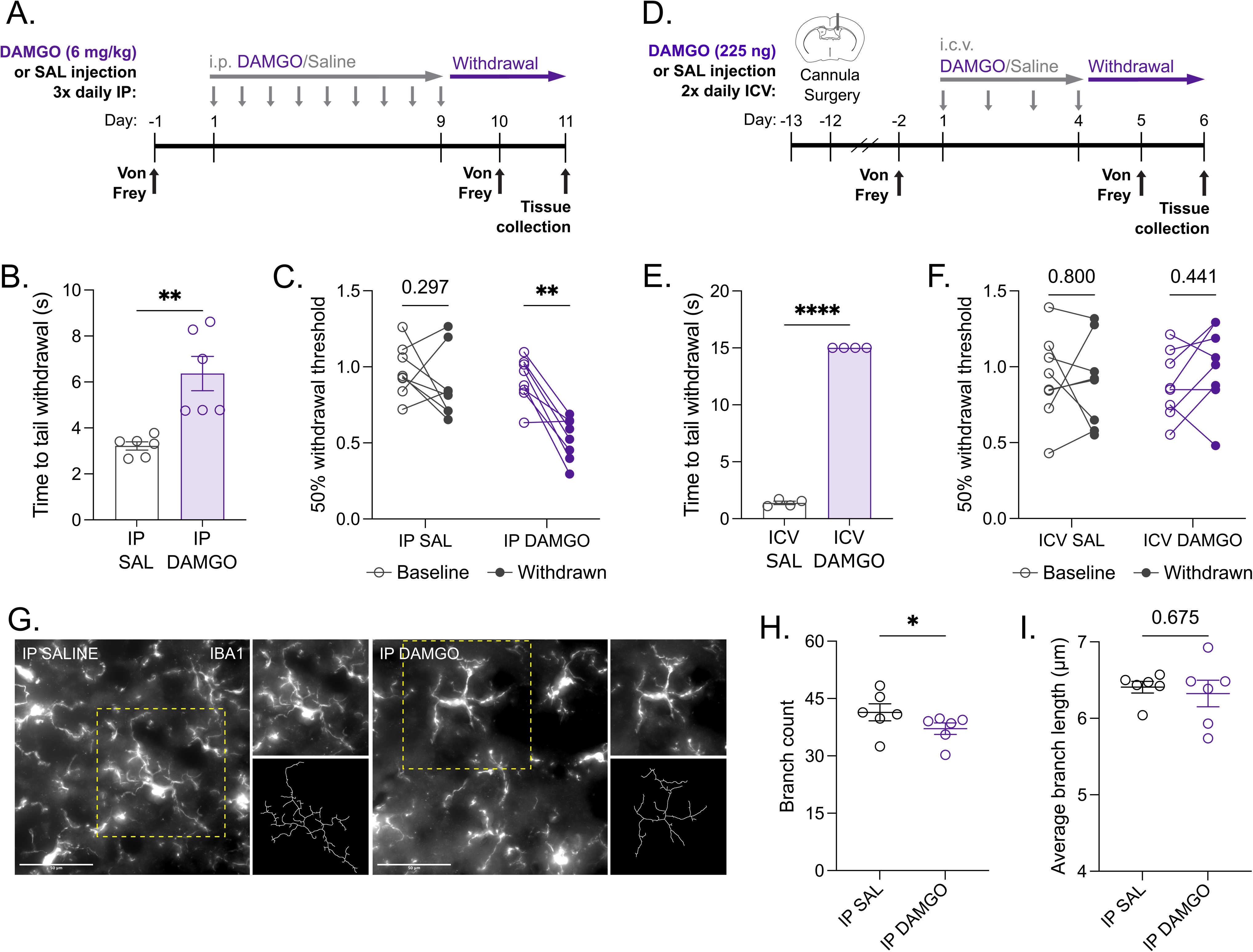
Peripheral, but not central, opioid exposure produces opioid-induced hyperalgesia. (**A**) Timeline of the experiment. (**B**) Acute analgesia following a single i.p. injection of 6 mg/kg DAMGO or saline (n_M_ = 6/group). (**C**) 50% paw withdrawal thresholds in mice treated with either i.p. DAMGO or saline for 9 days (n_M_ = 8/group). (**D**) Timeline of the experiment. (**E**) Acute analgesia following a single i.c.v. injection of 225 ng DAMGO or saline (n_M_ = 2, n_F_ = 2/group). (**F**) 50% paw withdrawal thresholds in mice treated with either i.c.v DAMGO or saline for 4 days (n_M_ = 4, n_F_ = 4/group). (**G**) IBA1^+^ microglia in the spinal dorsal horn of i.p. saline- or DAMGO-treated mice. (**H** and **I**) Branch counts (H) and average branch lengths (I) of skeletonized microglia from i.p. saline- or DAMGO-treated mice (n_M_ = 6/group). Data are represented as mean ± SEM; independent t test with Welch correction (B, E); Mann-Whitney U test (H, I); two-way repeated measures ANOVA with Holm-Sidak post-hoc (C, F). *p<0.05, **p<0.01, ****p<0.0001.

### Complete microglial depletion fails to alter opioid withdrawal in mice

To establish whether microglia were entirely dispensable for opioid-induced hyperalgesia, we used mice with a deletion of the Fms intronic regulatory element of the mouse *Csf1r* gene (*Fire*^Δ/Δ^ mice), which produces a complete and lifelong absence of microglia (56,57). In the lumbar spinal cord, *Fire*^Δ/Δ^ mice had no observable microglia (Fig. 4A,B), and, as previously described, highly diminished numbers of other (likely perivascular) macrophages compared to wildtype (WT) mice (60). (Fig. S6A-C). Loss of microglia did not alter the acute effects of opioids in mice, with comparable increases in locomotor activity following an acute 20 mg/kg morphine dose (Fig. 4C), and similar levels of acute morphine analgesia over a range of doses (Fig. 4D). Similarly, *Fire*^Δ/Δ^ mice also developed tolerance to the analgesic effects of morphine in the hotplate test after repeated administration (Fig. 4E, F), although this tolerance was slightly diminished in *Fire*^Δ/Δ^ males (Fig. S6D-F). In acute opioid withdrawal, *Fire*^Δ/Δ^ mice demonstrated comparable somatic withdrawal symptoms (including shaking/tremors, jumping, and writhing behaviour) to WT controls (Fig. 4G). Finally, the absence of microglia did not prevent the emergence of OIH in *Fire*^Δ/Δ^ mice, with opioid-withdrawn *Fire*^Δ/Δ^ mice demonstrating hypersensitivity in the coldplate (Fig. 4H), hotplate (Fig. 4I), tail-withdrawal (Fig. 4J), and Von Frey (Fig. 4K) assays.

**Figure 4.**
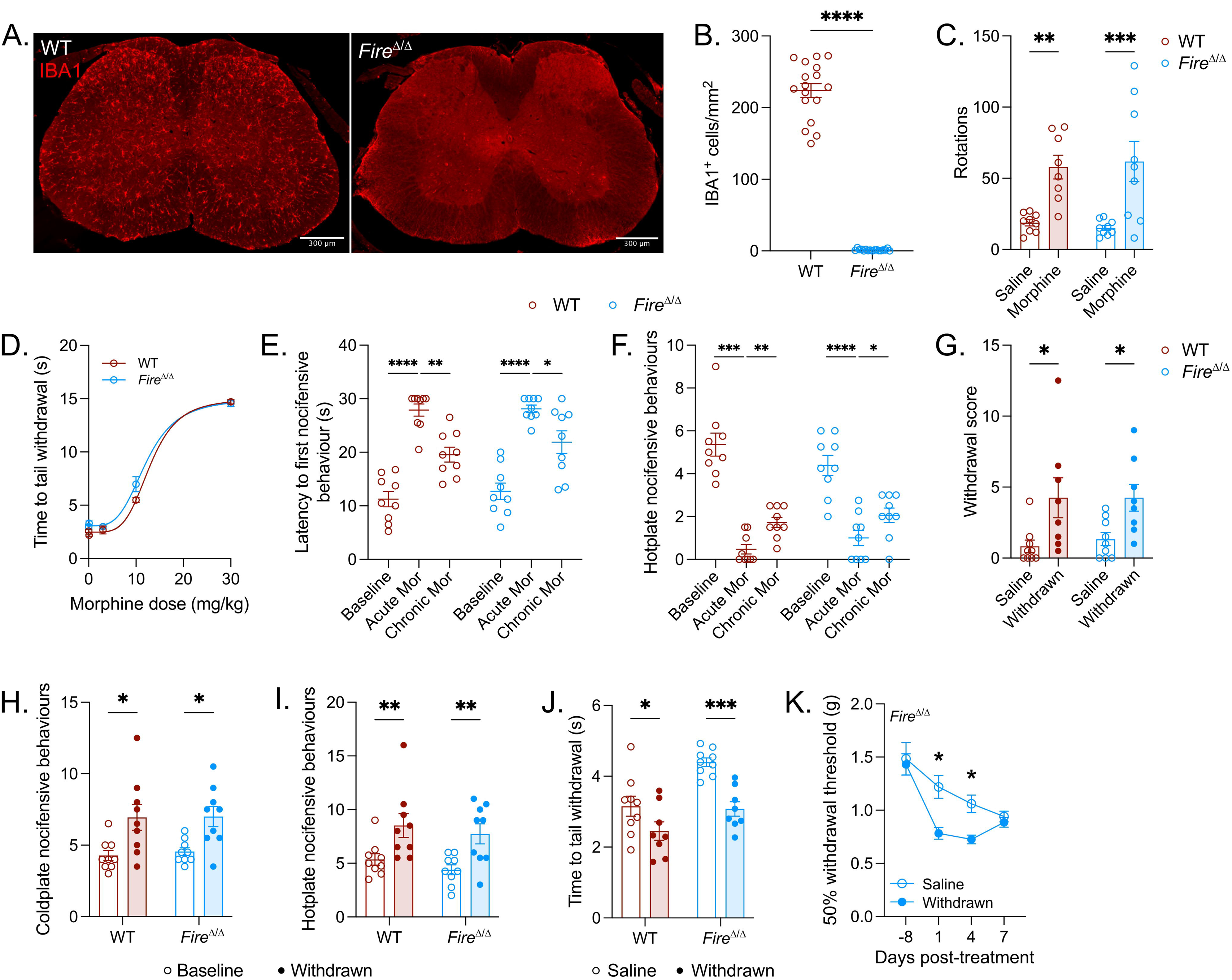
Microglial absence does not alter opioid withdrawal in mice. (**A**) Deletion of the Fms intronic regulatory element (*Fire*) super-enhancer of the *Csf1r* gene leads to lifelong absence of microglia in mice on a CBA/CaJ background (*Fire*^Δ/Δ^), scale bar = 300 µm. (**B**) IBA1^+^ cells in the spinal dorsal horn of wildtype (WT) and *Fire*^Δ/Δ^ mice (n_M_ = 3-4, n_F_ = 5/group). (**C**) Opioid-induced turning following injection of either saline or 20 mg/kg morphine in each genotype (n_M_ = 3-4, n_F_ = 5/group). (**D**) Morphine dose-response curves for WT and *Fire*^Δ/Δ^ mice (p = 0.099; n_M_ = 3-4, n_F_ = 5/group). (**E** and **F**) Latencies (E) and nocifensive behaviours (F) on a 53 °C hotplate at baseline and following a 10 mg/kg morphine injection in both opioid-naive (acute) and opioid-exposed (chronic) mice (n_M_ = 4, n_F_ = 5/group). (**G**) *Fire*^Δ/Δ^ mice display somatic withdrawal symptoms following prolonged opioid exposure (n_M_ = 3-4, n_F_ = 5/group). (**H** – **K**) During acute spontaneous withdrawal, opioid-exposed *Fire*^Δ/Δ^ mice show increased nocifensive behaviours in coldplate (H) and hotplate (I) assays (n_M_ = 4, n_F_ = 5/group), in addition to decreased tail withdrawal latencies (J) and reduced mechanical thresholds (K) (n_M_ = 3-4, n_F_ = 5/group). Data are mean ± SEM; Mann-Whitney U test (B); two-way ANOVA with Sidak-Holm post-hoc (C, G, J); two-way repeated measures ANOVA with Holm-Sidak post-hoc (E, F, H, I, K); non-linear regression with extra-sum-of-squares F test (D). *p<0.05, **p<0.01, ***p<0.001, ****p<0.0001.

## Discussion

In this study, we demonstrated that while spinal microglia display morphological changes during opioid withdrawal, these changes are accompanied by only modest transcriptional alterations. Despite their expression of the mu opioid receptor (MOR) transcript, *Oprm1*, direct exposure of microglia to opioids is dispensable for both morphological changes and the development of opioid-induced hyperalgesia (OIH) in mice. Finally, complete loss of microglia in *Fire*^Δ/Δ^ mice did not alter acute opioid analgesia, nor did it prevent the development of analgesic tolerance, somatic withdrawal, and OIH, arguing against microglial involvement in these phenomena.

Acute opioid analgesia is largely mediated through supraspinal actions (59–61), with a lesser contribution from spinal interneurons (24,62) and primary nociceptors (20,63–65). In contrast, the development of OIH largely appears to be a peripheral phenomenon, although modulation by supraspinal circuits in ongoing OIH is likely (25,26). Our finding that peripherally restricted opioid administration alone was able to induce mechanical hypersensitivity is in agreeance with reports that MOR activation on primary afferent nociceptors is necessary for maladaptive pro-nociceptive long-term potentiation of spinal cord circuitry (20–23). While microglia can modulate neuronal activity (66,67), given that *Fire*^Δ/Δ^ mice developed comparable OIH to their wildtype counterparts within the time frame examined, it is unlikely that this occurs to a significant degree in the context of OIH.

Previous work has shown that opioid administration in rodent models is associated with increases in the expression of myeloid-specific markers such as IBA1 and CD11b, which have been used has a correlate for increased glial reactivity. Similarly, others have demonstrated that opioid administration increases pro-inflammatory cytokine release associated with both opioid analgesic tolerance and hyperalgesia (29,31,38,68,69). We found no such increases in IBA1, CD11b, or classic pro-inflammatory cytokine expression in spinal microglia during withdrawal. Importantly, most studies have looked at changes in these markers during the period of opioid administration, instead of during withdrawal: in line with this, others have described apparent opioid-induced increases in proinflammatory cytokines that resolve within 24 hours of drug cessation (68). Consequently, while we cannot exclude the possibility that acute morphine exposure may elicit some degree of inflammation in spinal cord microglia, we have demonstrated that microglia do not adopt an inflammatory profile during either acute or prolonged opioid withdrawal, when hyperalgesia is most readily observable. However, given that bulk sequencing data of CD11b^+^ cells in the mouse spinal cord during chronic opioid administration has also revealed very little morphine-induced transcriptional alterations (70), it is unlikely that the microglial response to chronic opioid administration as compared to spontaneous withdrawal drastically differs.

In this study, we limited our analysis to spinal cord microglia, as this population has been most directly associated with OIH (30). Given both the differential impact of opioids across the brain and the region-specific variation in microglial states (13,71), we also cannot exclude the possibility that microglia in supraspinal regions may display a different withdrawal-associated phenotype than what we have described in this study. However, given that direct central administration of opioids failed to produce any evidence of OIH and *Fire*^Δ/Δ^ mice developed hyperalgesia despite total loss of microglia, it is unlikely that microglial-driven neuroinflammation in either spinal or supraspinal regions plays a significant role in the expression of OIH in rodents.

Notably, we observed alterations in microglial morphology at both 1- and 7-days post-drug, despite minor transcriptional alterations. Despite conflicting reports on the expression of opioid receptors by microglia (20,50), we observed *Oprm1* transcripts in spinal microglia, and others have described the expression of MOR protein by microglia in CX3CR1-eGFP-MOR-mCherry mice (51). cAMP dynamics in microglia are important regulators of morphology and process extension (72,73), and G_i_-coupled signaling in microglia results in process retraction (74). Consequently, it is possible that morphine alters microglial morphology directly via MOR-mediated suppression of cellular cAMP levels. On the other hand, peripherally-restricted opioid administration was sufficient to induce a modest but significant decrease in microglial processes, similar to what was observed with systemic administration, suggesting that direct opioid exposure may not be required for the morphological changes observed with morphine exposure. We also observed a small increase in the proportion of disease-associated microglia (DAM) in morphine-treated mice, which is a reactive transcriptional state associated with more amoeboid-like morphology (48). As our morphological measurements reflect an average of many individual cells, a small increase in the proportion of DAM-like microglia may account for the decrease in process number and total length that we observed here. In this study, we did not determine the mechanism through which opioids may drive an increase in this transcriptional state, although given the lack of response of microglia to opioids that we observed *in vitro* this is likely an indirect effect of chronic opioid administration. Ultimately, microglia are highly dynamic cells with an extremely diverse and extensive sensome, which enables them to react to a large variety of stimuli. However, the ability of these cells to detect shifts in the surrounding environment and initiate morphological changes in response does not necessarily implicate a functional outcome.

Multiple methods of microglial depletion have been employed previously, including use of CSF1R inhibitors and genetic manipulations involving diphtheria toxin. These methods, however, are often limited by the extent of depletion achieved, the specificity for microglia over other myeloid populations, and the variable susceptibility of different microglial states to depletion (75). PLX5622, in particular, interferes with the hepatic metabolism of multiple xenobiotics, including morphine (76), making it difficult to use in the context of opioid administration and withdrawal. In this study, we employed the use of *Fire*^Δ/Δ^ mice, who developed OIH and other withdrawal symptoms despite demonstrating a complete and total loss of microglia. In agreeance with our results, Liu et al. (47) also demonstrated that OIH develops in mice where microglia are depleted with PLX5622 administration or diphtheria toxin-mediated ablation, further supporting the idea that these cells are not necessary for the development of OIH. Additionally, we assessed the development of both analgesic tolerance and somatic withdrawal symptoms in *Fire*^Δ/Δ^ mice, and found that, like OIH, microglia are not required for the occurrence of these phenomena.

*Fire*^Δ/Δ^ mice, despite total loss of microglia, display minimal transcriptional alterations in neuronal and other glial cell types (77). While the fact that peripheral and not central opioid administration was sufficient to generate withdrawal-induced hypersensitivity makes it unlikely, we note that we cannot rule out a contribution of other glial cell types to the development of OIH (78,79), and a more thorough characterization of these cells during withdrawal would be of interest for future studies.

## Materials and Methods

### Animals

Male and female C57BL/6J mice aged 9-12 weeks at the beginning of each experiment were purchased from Jackson Laboratories (Bar Harbor, ME, cat #000640). For the detection of *Oprm1* transcripts in tdTomato^+^ microglia via in-situ hybridization, Cx3cr1*^CreER^* mice (The Jackson Laboratory, cat #021160) were crossed with ROSA26^tdT^ mice (The Jackson Laboratory, cat #007909) to produce *Cx3cr1*^CreER +/−^; ROSA26^tdT/-^offspring. *Cx3cr1*^CreER +/−^; ROSA26^tdT/-^pups were injected intraperitoneally with daily injections of 0.05 mL of 20 mg/mL tamoxifen (Sigma-Aldrich, cat #T5648) in corn oil (Sigma-Aldrich, cat #C8267) over postnatal days 13-15 to induce TdTomato expression in CX3CR1^+^ cells. Male and female *Csf1r*^ΔFIRE/ΔFIRE^ (*Fire*^Δ/Δ^) on a CBA/CaJ background (The Jackson Laboratory, cat #032783) and wildtype CBA/CaJ (The Jackson Laboratory, cat #000654) were purchased from Jackson Laboratories. Genotypes for the *Cx3cr1*^CreER +/−^; ROSA26^tdT/-^ and *Fire*^Δ/Δ^ mice were confirmed via real-time PCR (Transnetyx).

All animals were kept in a temperature- and humidity-controlled environment with free access to water and food. Mice were housed with littermates of the same sex in wire-top cages in groups of 2-5, except for cannulated mice, who were housed individually following surgery to prevent cannula removal by cagemates. Mice were kept on a 12 h:12 h dark/light cycle, with all behavioural experiments performed during the light cycle. Prior to all experiments, mice were habituated to behavioural testing rooms and handling. All animal experiments and procedures were conducted in compliance with the Canadian Council on Animal Care Guidelines and Policies with approval from the University of Alberta Health Sciences Animal Care and Use Committee.

### Drugs

Morphine sulfate pentahydrate powder (Toronto Research Chemicals) and [D-Ala2, N-Me-Phe4, Gly5-ol]-Enkephalin (DAMGO) acetate salt (Sigma-Aldrich, cat #E7384) were dissolved in 0.9% w/v NaCl for injection. Intraperitoneal (i.p.) injections were performed at a volume of 10 mL/kg. All control animals received an equivalent dose of saline. For all C57BL/6J mice, the morphine dosing regimen consisted of 4 days twice-daily escalating morphine injections (10, 20, 30, 40 mg/kg). For CBA/CaJ mice, the morphine dosing regimen consisted of 7 days of twice-daily escalating morphine injections (10, 20, 30, 40, 50, 50, 50 mg/kg).

### Behavioural testing

#### Measurement of locomotor activity

All animals received a habituation day to explore a rectangular arena (24 x 24.5 x 28 cm) for 10 min prior to the start of the experiment. Directly following i.p. injection of either morphine or saline control, mice were placed in the arena for 20 min, and the animal’s movement was tracked using video tracking software (Noldus Ethovision).

#### Von Frey test for mechanical allodynia

Mice were habituated to 7.5×10×7.5 cm plexiglass boxes suspended on a wire mesh platform. On test days, the up-down method (80,81) was used to determine mechanical nociceptive thresholds. Briefly, starting with a 0.4 g mechanical filament, filaments of various forces (0.04 g to 2 g) were applied to both hind paws in succession, following which negative and positive (withdrawing, flexing, and/or biting/licking of the hind paw) responses were recorded and used to calculate the 50% mechanical withdrawal threshold for each animal. Animals were tested by cage, and both the mesh platform and the plexiglass boxes were cleaned with ddH2O between cages. Experimenters were blinded to treatment during testing.

#### Thermal tail-withdrawal assay

Mice were habituated to the tail-withdrawal apparatus and restraint prior to the start of the experiment. On test days, mice were gently restrained, and approximately 2.5 cm of the distal portion of the tail was submerged in 49 °C water. The length of time the animal took to withdraw its tail after tail submersion was recorded, with a cut-off latency of 15 s established to prevent potential tissue damage. For assessing morphine analgesia, a single tail-withdrawal measurement was taken per animal at 20 min post-injection. For assessing baseline thermal thresholds and the development of opioid-induced hyperalgesia, 3 baseline measurements, separated by 20 min each, were taken for each animal per test, and withdrawal latencies across all 3 baselines were averaged. Experimenters were blinded to treatment during testing.

#### Dose-response curve

Morphine analgesic potency was determined using cumulative dose-response curves, with mice all receiving successive escalating doses of morphine (0.1, 3, 10, 30 mg/kg). Tail-withdrawal latencies were measured once every 20 min following each successive dose.

#### Thermal gradient test

For the thermal gradient test, mice were placed on a rectangular platform (110 cm x 18 cm) surrounded with black plastic walls (18.5 cm) and topped with a clear plexiglass cover. Cold and hot plates (Bioseb) were placed underneath either end of the platform, generating a thermal gradient from 2 °C to 65 °C along the long axis. All thermal gradient tests were performed in the dark, to minimize outside influences on place preference. Prior to the start of the experiment, mice received a habituation day, consisting of a 5 min exposure to the thermal gradient platform with the temperature off. On the test day, mice were placed on the platform with the hot/cold plates on and allowed to freely explore for 25 min. Overhead infrared cameras were used to record the position of the mouse along the platform, followed by subject tracking using Ethovision.

#### Hot and cold plate tests

For all hot and cold plate tests, mice were filmed after being placed on a thermal plate (Ugo Basile 35100) set to either 53 °C for the hot plate test, or 4 °C for the cold plate test. For the hot plate test, mice were scored for hind paw nocifensive responses, including licking/biting, shaking/lifting, and jumping, with a cut-off time of 30 s. For the cold plate test, mice were scored for both forepaw and hind paw nocifensive responses, including wringing and licking of the forepaw, and licking/biting, shaking, and flexing of the hind paw, with a cut-off time of 60 s. For all hot and cold plate tests, both the total number of nocifensive responses and the latency to first response was recorded: if an animal never exhibited a certain nocifensive behaviour, the latency was set as the cut-off time for the test. All videos were scored by two blinded experimenters, and the scores of each experimenter were averaged per animal.

#### Spontaneous withdrawal behaviour

For assessment of somatic withdrawal behaviour, mice were placed in a rectangular arena (see measurement of locomotor activity) on the first day of withdrawal post-morphine, and recorded for 20 min. The incidence of withdrawal-associated behaviour (shake/tremor, writhing, and jumping) was scored independently by two blinded experimenters, and the scores of each experimenter were averaged per animal.

### Intracerebroventricular (ICV) cannula implantation

Mice were anesthetized with 4% isoflurane before being placed in a stereotaxic frame on a 37 °C heating pad. Animals were maintained on 2% isoflurane throughout the surgery. 0.5% bupivacaine was injected intradermally at the incision site, and an incision was made to expose bregma and lambda. The skull was leveled using bregma as a reference, and manual craniotomy was performed using a 400 µm drill bit at the sites for cannula and bone screw insertion. 2 mm guide cannulas (Protech, cat #C315GS-5) were slowly inserted into the right lateral ventricle (A/P: -3.00, M/L: -1.00, D/L: -2.00), with two 1.6 mm bone screws (Protech, cat #00-96) placed posterior to the cannula insertion site. Inserted cannulas were fixed in place using dental cement (C&B Metabond) and covered using dummy cannulas with a 0.2 mm projection (Protech, cat #C315DCS-5). Immediately following surgery, mice were placed in heated cages for 1 hour, before being returned to their home cage and monitored daily for 3 days post-op alongside the ad-libitum administration of carprofen in drinking water. All mice were allowed to recover for at least 1 week before any experimental manipulations.

### ICV injections

For ICV injections, mice were gently restrained to immobilize the head, and dummy cannulas were removed from the guide cannula. Using a 10 µL Hamiliton syringe (Hamilton, cat #80300) attached to PE50 cannula tubing (Protech, cat #C313CT) and an internal cannula with a 0.45 mm projection (Protech, cat #C3151S-5), 2 µL of 112.5 ng/µL DAMGO (225 ng total dose) or sterile saline was administered into the right lateral ventricle at a rate of 1 µL/min. Internal cannulas were left in for 1 min post-injection, before being removed and replaced with the dummy cannula. All cannulated mice were injected with 2 µL Evan’s blue dye (Fisher Chemical, cat #E515) 90 min before euthanasia to confirm cannula placement.

### Microglia culture

#### Immunopanning and culture of primary mouse microglia

For primary microglial cultures, cortical tissue was dissected from postnatal day 5-7 CD1 mice (Charles River) and dissociated using the gentleMACs Neural Tissue Dissociation Kit P (Miltenyi Biotec cat #130–092-628). Dissociated tissue was pelleted through an ovomucoid (Worthington Biochemical) gradient, resuspended in a 0.02% bovine serum albumin (BSA; Sigma-Aldrich, cat #A4161) and DNase (Worthington cat #LS002007) solution, and filtered successively through 70, 50, and 20 µm cell strainers to remove debris. The resulting cell suspension was incubated in an anti-CD11b (clone M1/70; Invitrogen cat #14–0112-85) coated dish to select for microglia. Immunopanned cells were trypsinized (Sigma-Aldrich, cat #T9935) for 4 min at 37 °C to decrease adherence to the panning dish and dislodged by pipetting in ice-cold media. Dislodged cells were counted and plated at a density of 20,000 microglia per well. Culture surfaces were coated in poly-d-lysine (10 mg/mL; Sigma-Aldrich cat #P6407) to promote cell adherence. Microglia were grown in serum-free media, containing Neurobasal (Gibco, cat #21103049), 1X Gem21 (GeminiBio, cat #400–160-010) and N2 (ThermoFisher, cat #17502–048) supplements, 2 mM GlutaMax (Gibco, cat #35050–061), 1 mM sodium pyruvate (Gibco, cat #11360–070), penicillin-streptomycin (Gibco, cat #15140122), 1 mg/mL lipated BSA (ThermoFisher, cat #11020021), 50 mM sodium chloride (Sigma-Aldrich, cat #S5150), 10 ng/mL d-biotin (Sigma-Aldrich, cat #B4639), and 0.2% lactic acid (Sigma-Aldrich, cat #L1250). To facilitate survival in serum-free conditions, microglial growth media was supplemented with 1.5 µg/mL cholesterol (Avanti Polar Lipids, cat #700000P), 100 ng/mL oleic acid (Cayman, cat #90260), 1 ng/mL gondoic acid (Cayman), 1 µg/mL heparin sulfate (Galen Laboratory Supplies, cat #20606), 10 ng/mL CSF-1 (Peprotech, cat #315–02), and 2 ng/mL TGF-b (Peprotech, cat #100-35B). Microglia were cultured for 5-7 days in serum-free media before all experiments.

#### Phagocytosis assay in primary microglia

For live-cell imaging experiments to track phagocytosis in cultured microglia, microglia were pre-labelled with CellTracker Red (1 µM; ThermoFisher, cat #C34552) and NucBlue (10 drops/mL; ThermoFisher, cat #R37605). Cells were then treated with morphine sulfate (Toronto Research Chemicals) or PBS vehicle alongside the addition of pHrodo-labeled (Invitrogen, cat #P36600) myelin (2 µg/mL) or nuclei (5000/well). For experiments using nuclei, cells were imaged before the addition of nuclei to ensure that isolated nuclei were not counted towards the total microglial cell density in each well. Following treatment, cells were immediately imaged at 3-hour intervals over the course of 24 hours at 10X (ImageXpress Pico; MolecularDevices). Image analysis was performed in CellReporterXpress using the wavelength cell-scoring analysis function; output was filtered by cell and nuclei size to exclude debris, and images were visually examined to check the accuracy of the automated counting.

#### Viability assay in primary microglia

To assess microglial viability in drug treatments, primary mouse microglia were incubated in various morphine concentrations or PBS control for 24 hours at 37 °C, following which the treatment-containing media was removed and replaced with media containing NucBlue (10 drops/mL), 2 µg/mL Calcein AM (Invitrogen, cat #C1430), and 4 µg/mL propidium iodide (PI; Invitrogen, cat #P3566) in Neurobasal base. Cells were then incubated at 37 °C for 10 min, before being imaged at 10X and analyzed as described for the phagocytosis assay. Cells were scored as live (Calcein AM^+^PI^-^), dead (PI^+^), or unlabeled (Calcein AM^-^PI^-^).

### Immunohistochemistry

Mice were anesthetized with an i.p. injection of 100 mg/kg sodium pentobarbital (Euthanyl, Western Drug Distribution Centre Ltd, DIN #00141704) and perfused transcardially with 10 mL 0.9% saline, followed by 4% paraformaldehyde (PFA). Spinal cords were dissected and fixed at 4 °C in 4% PFA for 24-48 hr. Fixed spinal cords were cryoprotected by incubating in 30% sucrose at 4 °C, before being frozen in Optimal Temperature Cutting Compound (OCT; TissueTek, cat #4585) for cryosectioning. The lumbar region of the spinal cord was sectioned at 20 µm (Leica CM3050S) and mounted on Superfost PlusTM slides (Fisher Scientific, cat #1255015); slides were stored at -80 °C until use. Tissue was blocked for 1 hr with 10% normal donkey serum (NDS; Sigma-Aldrich, cat #D9663) in 0.1M PBS containing 0.2% Triton-X100 (Sigma-Aldrich, cat #X100). Sections were incubated overnight at 4 °C in rabbit IBA1 (1:500; Wako, cat #019-19741) or rat CD45 (1:100; BD Pharmingen, cat #550539). All antibody solutions were prepared 0.2% Triton-X100 containing 2% NDS and 2% bovine serum albumin (BSA; Sigma-Aldrich, cat #A4161). The next day, sections were labelled with the appropriate secondary: either donkey anti-rabbit Alexa Fluor 488 (Invitrogen, cat #A21206) or donkey anti-rat Alexa Fluor 647 (Invitrogen, cat #A78947) at 1:200. Images were obtained on a Zeiss AxioImager microscope (Carl Zeiss, Oberkochen, Germany) at either 20X for whole dorsal horn image analysis, or 63X for skeleton analysis. All image analysis was performed using ImageJ (NIH). For analysis of microglial morphology, microglia were thresholded by IBA1 expression, and binary images were skeletonized and analyzed using the Analyze Skeleton plugin.

### Fluorescent in situ hybridization

In situ hybridization was performed using the RNAscope multiplex fluorescent reagent kit v2 (Advanced Cell Diagnostics (ACD), California, USA, cat #323100). *Cx3cr1*^CreER +/−^; ROSA26^tdT/-^mice were perfused with PBS followed by 4% PFA prior to spinal cord dissection. Spinal cords were frozen in OCT, sectioned at 20 µm, and mounted on to Superfrost PlusTM slides. RNAscope was performed using the standard RNAscope multiplex protocol: briefly, slides were baked at 60 °C for 30 min, post-fixed in 4% PFA for 15 min, and then dehydrated in ethanol. Dehydrated slides were treated with H_2_O_2_ for 10 min before being boiled in target retrieval reagent for 15 min and permeabilized with protease III for 30 min at 40 °C. Samples were hybridized to the *Oprm1* probe (Mm-Oprm1; ACD #315841) for 2 hr at 40 °C, before the signal was amplified with amplifiers 1-3 and labelled with Opal 690 (Akoya Biosciences, #FP1497001KT). All samples were run alongside both negative (RNAscope negative control probe-*Dapb*; ACD, #310043) and positive (RNAscope 3-plex positive control probe-Mm; ACD, #320881) controls. Immediately following the RNAscope assay, slides were blocked with 10% NDS for 1 hr, before being incubated overnight at 4 °C with goat anti-TdTomato (1:200; Sicgen, cat #AB8181). Slides were labelled with donkey anti-goat Alexa Fluor 555 (Invitrogen, cat #A21432) for 45 min, and coverslipped with ProLong Gold antifade mountant containing DAPI (ThermoFisher, cat #P36931). Slides were imaged using a Zeiss AxioImager microscope with a 40X objective lens. Cells were counted as *Oprm1*^+^ if they had ≥3 *Orpm1* puncta within the cell soma.

### RNA extraction and quantitative RT-PCR

For the extraction of total RNA from mouse spinal cords, mice were anesthetized with sodium pentobarbital (as above) and perfused transcardially with 10 mL 0.9% saline, followed by dissection of the lumbar spinal cord. Spinal cords were homogenized and immediately flash-frozen in QIAzol tissue lysis reagent (Qiagen, cat. #79306), before being stored at -80 °C for later processing. Crude RNA was extracted using a phenol:chloroform extraction, and processed with RNeasy micro plus kits (Qiagen, cat #74034). 500 ng of total RNA was processed with DNase I (Invitrogen, cat #18068-015) and primed with oligo(dT)12-18 (Invitrogen, cat #18418-012) before being reverse-transcribed using SuperScript III (Invitrogen, cat #56575). qPCR was run using Fast SYBR Green master mix (Applied Biosystems, cat #4385612) on a QuantStudio3 real-time PCR instrument (Applied Biosystems). Gene expression was normalized over the mean of two reference genes (*Rpl5* and *Ppia*). Qiagen RT2 qPCR primers were used for all qPCR measurements, including *Rpl5* (cat #PPM25102A), *Ppia* (cat #PPM03717B), *Itgam* (cat #PPM03671F), *Il1b* (cat #PPM03109F), *Il6* (cat #PPM03015A), and *Tnf* (cat #PPM03113G).

### Cell isolation

Following anesthetization with sodium pentobarbital and transcardial perfusion with 20 mL 0.9% saline, lumbar spinal cords were dissected and immediately placed in cold Hank’s Buffered Salt Solution (HBSS; Gibco, cat #14175-079). For the isolation of cells for single cell sequencing, spinal cords from mice within each treatment group were processed together via cold mechanical dissociation, where spinal cords were finely minced, and homogenized in Dulbecco’s Modified Eagle Medium (DMEM; Gibco, cat #11960-044) supplemented with 10 µg/mL DNase (Worthington cat #LS002007) in a dounce homogenizer with a loose-fitting pestle on ice. For the isolation of cells for flow cytometry analysis, spinal cords from individual mice were minced and incubated in an enzyme digestion mix containing 1 mg/mL collagenase D (Roche, cat #11088858001) and 50 µg/mL DNase in RPMI 1640 (Gibco; cat # 11875093) supplemented with 2% fetal bovine serum (FBS). Samples were incubated in the digestion mix for 30 min at 37 °C, with gentle manual dissociation with an 18G needle after 15 min. Following either mechanical or enzymatic dissociation, samples were filtered successively with 100 µm and 70 µm cell strainers, followed by washing with DMEM containing 10 µg/mL DNase. Samples were centrifuged (300xg, 10 min, 4 °C) and resuspended in HBSS, before being gently mixed with debris removal solution (Miltenyi Biotec, cat #130-109-398), and overlaid with HBSS. Samples were centrifuged (3000xg, 10 min, 4 °C), and following aspiration of the top two layers, were topped up to 14 mL with HBSS and gently inverted. Cells were pelleted (1000xg, 10 min, 4 °C) and resuspended in HBSS before being filtered through a 20 µm cell strainer.

### Flow cytometry

Following cell isolation, samples were incubated with Zombie Aqua fixable viability kit (1:200 in HBSS; BioLegend, cat #423102) for 15 min at 4 °C, washed with HBSS before centrifugation (300 xg, 5 min, 4 °C), and resuspended in anti-mouse CD16/32 (1:200; eBiosciences, cat #14-0161-82) in HBSS supplemented with 1% BSA for 10 min at 4 °C. Cells were stained in the blocking solution with PE/Cyanine5 CD45 (1:200; BioLegend, cat #103110), FITC CD11b (1:200; BioLegend, cat #101205), PE Ly-6G/Ly-6C (1:100; Gr-1, BioLegend, cat #108407), and APC/Fire™ 750 Ly-6C (1:200; BioLegend, cat #128045) for 30 min at 4 °C. Single-stained controls were prepared for all samples using cells from naïve mice, apart from the use of compensation beads (BD Biosciences, cat #552844) for PE-Gr1 and APC/Fire^TM^ 750-Ly6C. Following a wash with HBSS, cells were fixed with 2% PFA for 15 min at 4 °C and used for flow cytometric analysis on a spectral flow cytometer (Cytek Northern Lights) at the University of Alberta Flow Cytometry Core. Data was unmixed using SpectroFlo (v4.0.3) and analyzed using FlowJo (v.10.9.0). Microglia were analyzed by gating for time, single cells, and live cells (Zombie Aqua negative), and defined as CD45^+^CD11b^+^Gr1^-^.

### Library preparation and sequencing

Single-cell sequencing experiments were done in two independent batches, with each batch containing a sample from each of the three conditions, for a total of two replicates per condition. Each sample contained the pooled lumbar spinal cords of either 6 (batch 1) or 8 (batch 2) male mice from each treatment group, and samples within each batch were processed and sequenced concurrently. Following cold mechanical cell isolation as described above, cells were incubated in anti-mouse CD16/32 (1:200) in HBSS supplemented with 1% BSA for 5 min at RT and then incubated in Zombie Aqua fixable viability kit (1:200), DRAQ5 fluorescent probe solution (1:1000; ThermoFisher, cat #62254), and PE CD11b (1:200; BioLegend, cat #101208). Live, nucleated, CD11b^+^ cells were sorted on either a Sony MA900 (batch 1) or a BD FACSAria III (batch 2) at the University of Alberta Flow Cytometry Core, and manually counted with a hemocytometer prior to library prep. Sorted cells were processed using the 10x Genomics Chromium Next GEM Single Cell 3’ Gene Expression Dual Index v3.1 kit (10x Genomics, cat #PN-1000269) and sequenced on an Illumina NovaSeq X Plus PE150 (Novogene). FASTQ files were aligned to the GRCm39-2024-A mus musculus reference genome using the cellranger count pipeline (v.9.0.1).

### Single-cell transcriptomics

#### Quality control and clustering

Transcriptomic analysis, including quality control, subsetting, clustering, and differential expression (DE) were performed using Seurat (v5.5.0) in an R statistical environment (v4.5.0). Each sample was processed with SoupX (v1.6.2) to remove ambient RNA contamination, followed by initial filtering of the data to include only genes expressed in a minimum of 3 cells and cells with a minimum of 300 features and 1500 RNA transcripts, and a maximum of 5% mitochondrial reads. To remove doublets, scDblFinder (v1.22.0) was used to label and exclude presumptive doublets, and cells with total UMI counts in the top 0.5% for each batch were excluded. Each sample was normalized using the SCTransform (3000 highly variable genes) function, and samples were integrated using Seurat’s native reciprocal PCA (RPCA) integration. Clustering using the first 30 principal components was performed to identify cell types, and microglial clusters (defined by high expression of *Tmem119* and *P2ry12*) were subset from other CD11b^+^ cell populations and further filtered with cell type specific thresholds (>1200 total features and <4% mitochondrial reads). Cells expressing high levels of marker genes for non-microglial cells were manually removed. The microglial subset was re-normalized with SCTransform (1000 highly variable features and mitochondrial read percentage regressed) and integrated by batch using Harmony (v2.0.5). One gene (*Kazn*) was manually removed from the highly variable genes list, as its inclusion produced a small cluster defined by this single gene, with no other differentially expressed genes or association with quality control metrics. An elbow plot was used to select the number of principal components (1–18) with which to run UMAP and clustering. Clustering resolution was selected based on visualization of a clustering tree spanning a range of resolutions generated by the clustree (v0.5.1) package and further refined using the Single Cell Clustering Assessment Framework (SCCAF; v0.0.10) package with a minimum accuracy of 90% in a Jupyter Notebook (v1.1.1) running a python environment (v3.9.23). To generate wiggle plots with reads mapped to the genomic locus containing *Oprm1*, BAM files for each sample were subset to include only cells identified as microglia in the single-cell data analysis using subset-bam (v1.1.0) and visualized in the Integrative Genomics Viewer (IGV) (81).

#### Differential expression (DE) and over-representation analysis (ORA)

DE genes were determined in Seurat using a Wilcoxon rank-sum test via the FindMarkers or FindAllMarkers functions with a log fold change threshold of 0.2 and a minimum expression of either 15% (for cluster comparisons) or 20% (for treatment comparisons). DE genes with adjusted p values < 0.05 were used for ORA using the R package clusterProfiler (v4.16.0). For pseudobulk comparisons, Seurat’s AggregateExpression function was used to aggregate summed RNA counts per sample, and a Wald test (DESeq2 v1.48.2) was used to identify DE genes with batch as a covariate. For the scoring of the DAM gene module, individual cells were scored on their expression of DAM-associated genes (50) using the AddModuleScore function, and groups were compared using independent t tests with a Bonferroni post-hoc correction.

### General statistical analysis

Statistical analysis was performed using GraphPad Prism software version 11.0.2. All data was tested for normality, and parametric or non-parametric statistics were used accordingly. For comparisons of two groups, independent t tests or Mann Whitney U tests were performed, while the means of >2 groups were analyzed using 2- or 3-way analysis of variance (ANOVA) followed by Sidak-Holm post-hoc analysis to correct for multiple comparisons. All data is represented as mean ± SEM, and the details of statistical testing, as well as the number of male (n_M_) and female (n_F_) mice used, are included in the figure captions. For all experiments, an alpha value of 0.05 was set.

## Acknowledgments

We would like to thank the Faculty of Medicine and Dentistry Flow Cytometry Core at the University of Alberta for assistance with cell sorting and analysis.

## Author contributions

J.E.R.N and A.M.W.T designed experiments. J.E.R.N performed most of the experiments and analyzed all data. K.V.L, T.W., J.M.C, A.C.W, C.S.B, M.F.S.H, K.M.S, B.S, and M.E.T performed some experiments. J.E.R.N wrote the manuscript, with proofreading by A.M.W.T and J.R.P.

## Data and code availability

All data and code reported in this paper will be shared by the lead contact upon request.

Single-cell sequencing data will be deposited in the GEO database and made publicly available as of the date of publication.

## Figure legends

**Fig. S1. Opioid withdrawal does not produce significant activation of spinal microglia.**

**(A**) Gating strategy for spinal cord microglia, following gating for time, single cells, and live cells.

(**B**) The proportion of spinal cord microglia is not altered at day 4 post-treatment in saline-treated and morphine-withdrawn mice (n_M_ = 2, n_F_ = 3/group).

(**C**) Median fluorescent intensity (MFI) of CD45 and CD11b in sorted microglia at day 4 post-treatment (n_M_ = 2, n_F_ = 3/group).

(**D** - **E**) Integrated density of IBA1 (D) and CD45 (E) in spinal dorsal horn microglia at day 1 post-treatment (n_M_ = 3-4, n_F_ = 2/group).

(**F**) Expression of Itgam (CD11b) in the lumbar spinal cord of saline-treated and morphine-withdrawn mice at days 1 and 7 post-treatment (day 1: n_M_ = 4, n_F_ = 2/group; day 7: n_M_ = 2, n_F_ = 4/group).

(**G**) Expression of *Il1b*, *Il6*, and *Tnf* are unchanged in the lumbar spinal cord at day 1 post-treatment (n_M_ = 4, n_F_ = 2/group).

Data is represented as mean ± SEM, independent t test with Welch correction (B, D, E); multiple independent t tests with Sidak-Holm correction (C, F, G).

**Fig. S2. Quality control following scRNA-sequencing of mouse spinal cords.**

(**A** and **B**) Feature counts and unique molecular identifier (UMI) counts across samples (A), and percent mitochondrial genes and UMI counts across samples (B).

(**C** - **E**) Total UMI counts, total feature counts, and percent mitochondrial genes across each treatment group.

(**F**- **H**) Total UMI counts, total feature counts, and percent mitochondrial genes across each microglial cell cluster.

(**I** and **J**) Percent mitochondrial genes (I) and total feature counts (J) per cell represented on a UMAP of all microglial cells.

**Fig. S3. Extended scRNA-sequencing and pseudobulk testing.**

(**A**) Representative wiggle plots of microglial reads mapped to the mouse *Oprm1* gene. Exons (yellow) present in various splice variants are numbered below.

(**B**) Proportions of microglia in each cluster within each of the three treatment groups.

(**C** and **D**) Volcano plots of day 1 (C) and day 7 (D) differentially expressed genes following DESeq2 on pseudobulked counts (n = 2 replicates/group).

**Fig. S4. Morphine does not alter microglial phagocytosis *in vitro*.**

(**A**) Representative 10x image of primary mouse microglia labelled with DAPI and CellTracker Red, and incubated for 24 hr with phrodo-labelled myelin.
(**B**) % fold change in pHrodo^+^ cells after incubating cells with pHrodo-labelled myelin for 24 hr with PBS or various concentrations of morphine. Values are normalized to the PBS average within each independent experiment: triangles represent the mean of replicates (n = 2) and circles represent individual wells (n = 3) within each replicate; replicates are denoted by alpha value.
(**C**) % phrodo^+^ cells after 24 hr incubation in pHrodo-labelled nuclei with either PBS or 10 µM morphine (n = 3 wells in 1 independent experiment).
(**D**) Morphine did not alter the proportion of live (Calcein AM^+^ PI^-^) cells following a 24 hr incubation (average of 2 independent replicates, n = 3 wells/replicate).

Data is represented as mean ± SEM, one-way Brown-Forsythe and Welch ANOVA (B); unpaired t test (C).

**Fig. S5. Extended data for mice treated with i.p. or i.c.v. DAMGO**

(**A**) Distance travelled in an open field arena after a single i.p. injection of saline or 6 mg/kg DAMGO (n_M_ = 3/group).

(**B** - **D**) Morphology of spinal dorsal horn microglia in mice chronically treated with i.p. saline/DAMGO (n_M_ = 6/group)

(**E** -**I**) Morphology of spinal dorsal horn microglia in mice chronically treated with i.c.v. saline/DAMGO (n_M_ = 4, n_F_ = 4/group).

Data are represented as mean ± SEM; independent t test (A); Mann-Whitney U test (B-I). *p<0.05.

**Fig. S6. Extended data for *Fire*^Δ/Δ^ mice**

(**A**) Spinal dorsal horn images at 20X in WT and *Fire*^Δ/Δ^ mice, scale bar = 100 µm.

(**B** and **C**) Perivascular macrophages (IBA1^+^ CD45^hi^) in the spinal dorsal horn are decreased in *Fire*^Δ/Δ^ mice, while IBA1^-^CD45^hi^ cell densities remain unchanged (C) (n_M_ = 3-4, n_F_ = 5/group).

(**D** – **F**) Hotplate latencies to first nocifensive behaviour (D), hotplate nocifensive behaviours (E), and tail withdrawal latencies (F) following a 10 mg/kg acute morphine challenge in WT and *Fire*^Δ/Δ^ mice treated chronically with morphine, separated by sex (n = 3-5/group).

Data are mean ± SEM; independent t test with Welch correction (B, C); two-way ANOVA with Holm-Sidak post-hoc (F); three-way repeated measures ANOVA (D, E). *p<0.05, **p<0.01, ****p<0.0001.

