## Supplemental Files for "Spinal microglia are dispensable for the development of opioid-induced hyperalgesia"

### Figure S1

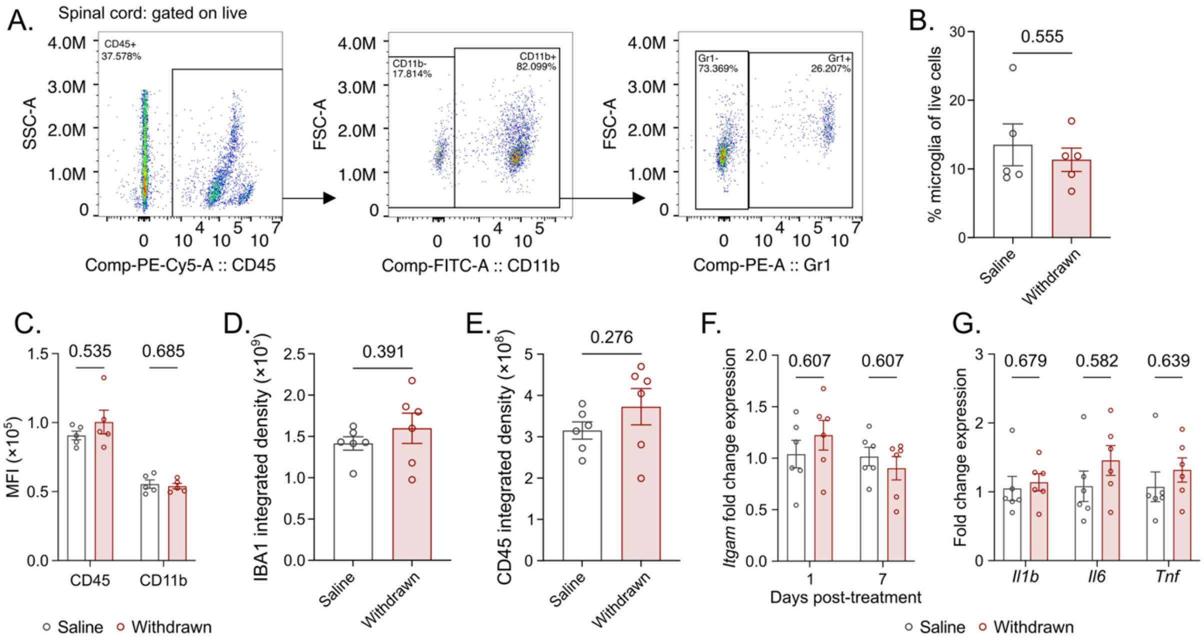


**Fig. S1. Opioid withdrawal does not produce significant activation of spinal microglia.**

(**A**) Gating strategy for spinal cord microglia, following gating for time, single cells, and live cells.

(**B**) The proportion of spinal cord microglia is not altered at day 4 post-treatment in saline-treated and morphine-withdrawn mice ($n_M = 2$, $n_F = 3$/group).

(**C**) Median fluorescent intensity (MFI) of CD45 and CD11b in sorted microglia at day 4 post-treatment ($n_M = 2$, $n_F = 3$/group).

(**D-E**) Integrated density of IBA1 (D) and CD45 (E) in spinal dorsal horn microglia at day 1 post-treatment ($n_M = 3$-4, $n_F = 2$/group).

(**F**) Expression of Itgam (CD11b) in the lumbar spinal cord of saline-treated and morphine-withdrawn mice at days 1 and 7 post-treatment (day 1: $n_M = 4$, $n_F = 2$/group; day 7: $n_M = 2$, $n_F = 4$/group).

(**G**) Expression of *Il1b*, *Il6*, and *Tnf* are unchanged in the lumbar spinal cord at day 1 post-treatment ($n_M = 4$, $n_F = 2$/group).

Data is represented as mean ± SEM, independent t test with Welch correction (B, D, E); multiple independent t tests with Sidak-Holm correction (C, F, G).

Figure S2

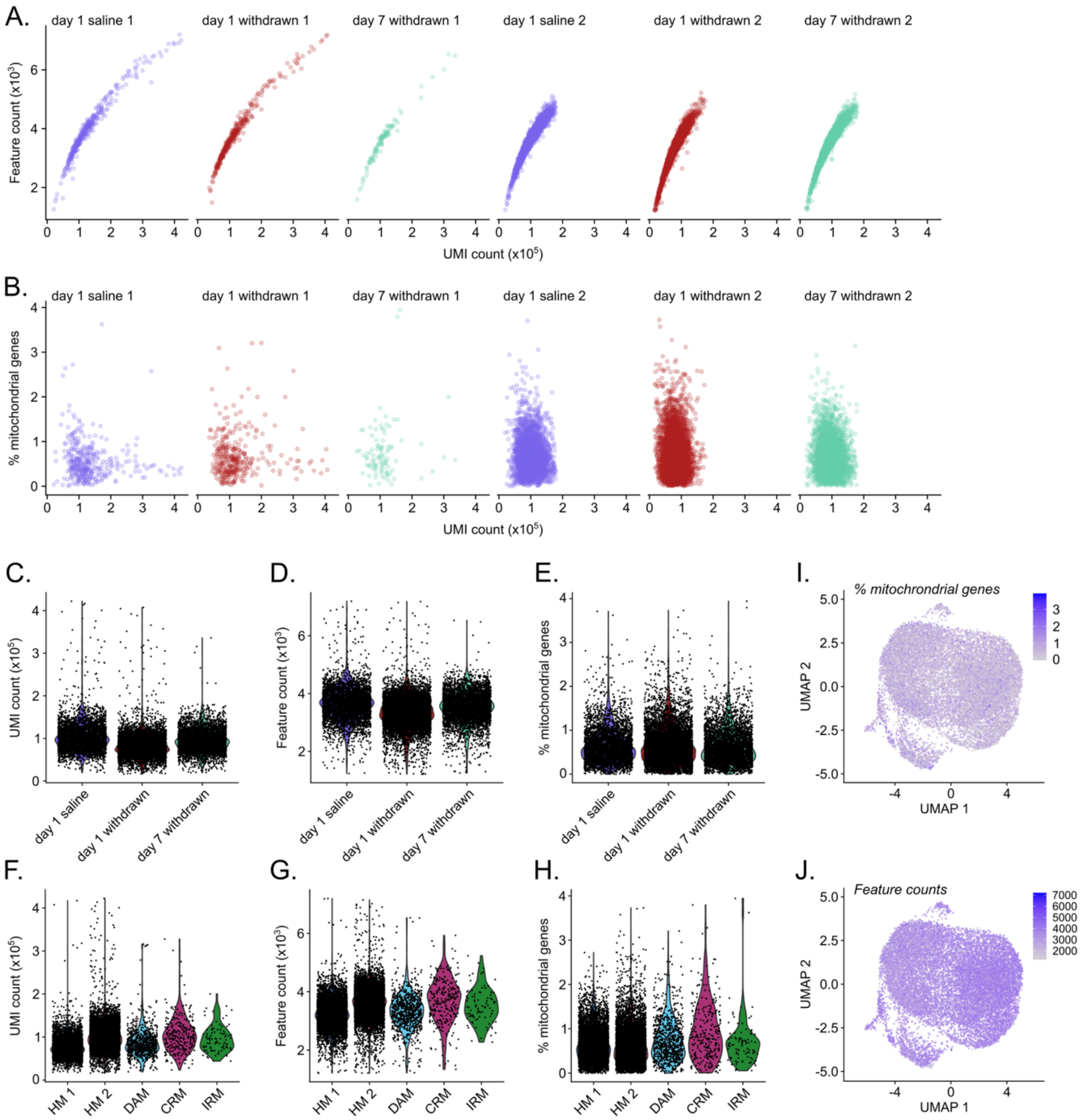


**Fig. S2. Quality control following scRNA-sequencing of mouse spinal cords.**

(**A** and **B**) Feature counts and unique molecular identifier (UMI) counts across samples (A), and percent mitochondrial genes and UMI counts across samples (B).

(**C** - **E**) Total UMI counts, total feature counts, and percent mitochondrial genes across each treatment group.

(**F**- **H**) Total UMI counts, total feature counts, and percent mitochondrial genes across each microglial cell cluster.

(**I** and **J**) Percent mitochondrial genes (I) and total feature counts (J) per cell represented on a UMAP of all microglial cells.

Figure S3

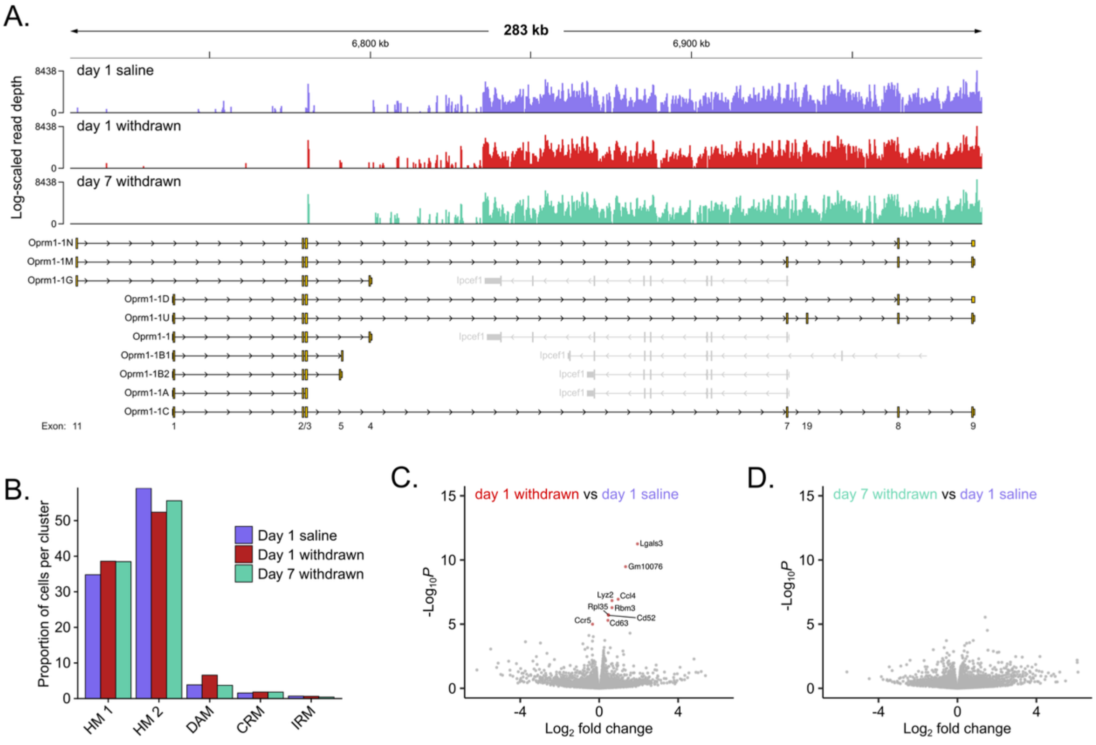


**Fig. S3. Extended scRNA-sequencing and pseudobulk testing.**

(**A**) Representative wiggle plots of microglial reads mapped to the mouse *Oprm1* gene. Exons (yellow) present in various splice variants are numbered below.

(**B**) Proportions of microglia in each cluster within each of the three treatment groups.

(**C** and **D**) Volcano plots of day 1 (C) and day 7 (D) differentially expressed genes following DESeq2 on pseudobulked counts (n = 2 replicates/group).

Figure S4

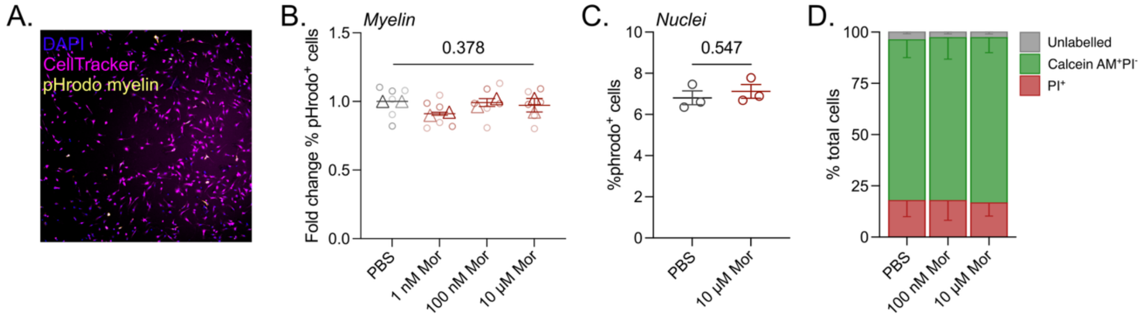


**Fig. S4. Morphine does not increase microglial phagocytosis *in vitro*.**

(**A**) Representative 10x image of primary mouse microglia labelled with DAPI and CellTracker Red, and incubated for 24 hr with phrodo-labelled myelin.
(**B**) % fold change in pHrodo$^{+}$ cells after incubating cells with pHrodo-labelled myelin for 24 hr with PBS or various concentrations of morphine. Values are normalized to the PBS average within each independent experiment: triangles represent the mean of replicates (n = 2) and circles represent individual wells (n = 3) within each replicate; replicates are denoted by alpha value.
(**C**) % phrodo$^{+}$ cells after 24 hr incubation in pHrodo-labelled nuclei with either PBS or 10 µM morphine (n = 3 wells in 1 independent experiment).
(**D**) Morphine did not alter the proportion of live (Calcein AM$^{+}$ PI$^{-}$) cells following a 24 hr incubation (average of 2 independent replicates, n = 3 wells/replicate).
Data is represented as mean ± SEM, one-way Brown-Forsythe and Welch ANOVA (B); unpaired t test (C).

### Figure S5

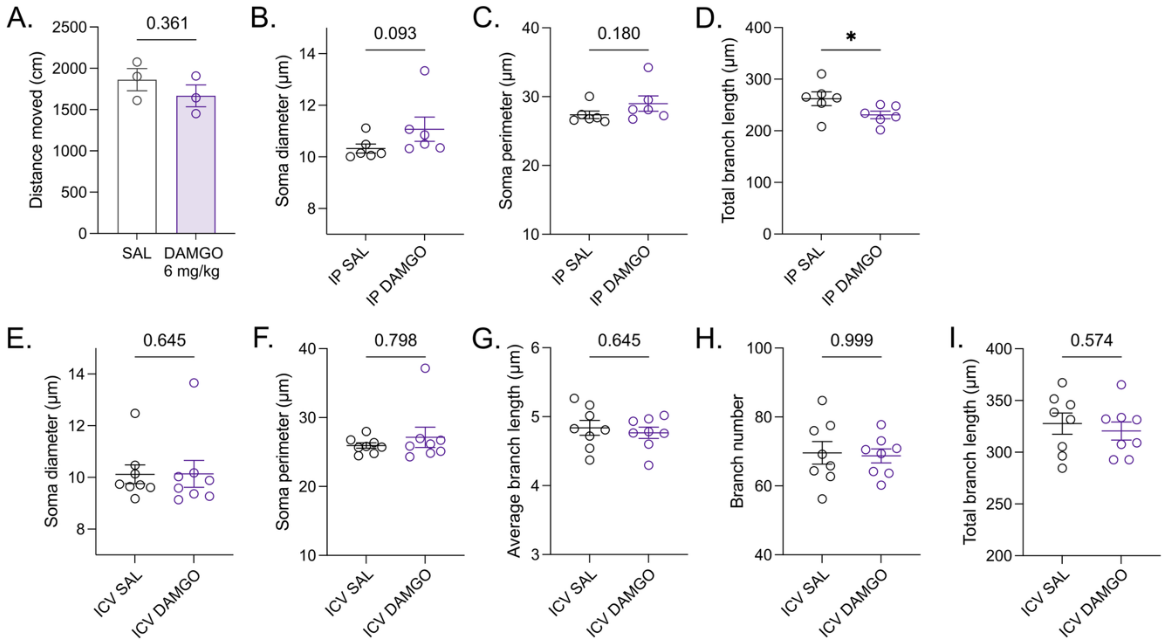


**Fig. S5. Extended data for mice treated with i.p. or i.c.v. DAMGO**

(**A**) Distance travelled in an open field arena after a single i.p. injection of saline or 6 mg/kg DAMGO ($n_M$ = 3/group).

(**B** - **D**) Morphology of spinal dorsal horn microglia in mice chronically treated with i.p. saline/DAMGO ($n_M$ = 6/group)

(**E** -**I**) Morphology of spinal dorsal horn microglia in mice chronically treated with i.c.v. saline/DAMGO (E-I) ($n_M$ = 4, $n_F$ = 4/group).

Data are represented as mean ± SEM; independent t test (A); Mann-Whitney U test (B-I). *$p<0.05$.

### Figure S6

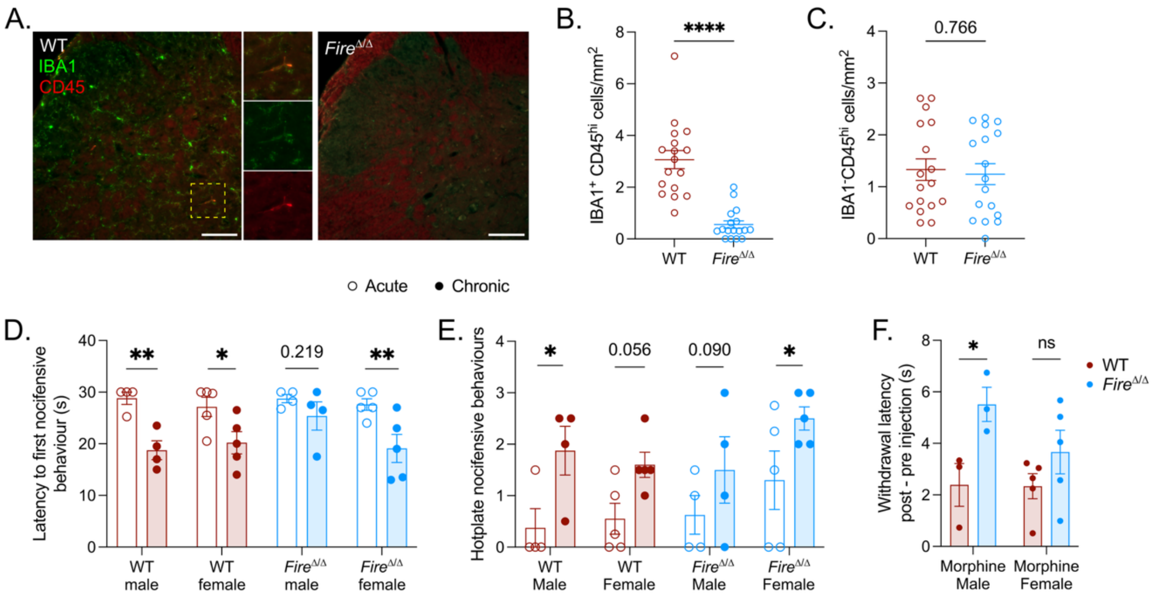


**Fig. S6. Extended data for *Fire*$^{\Delta/\Delta}$ mice**

(**A**) Spinal dorsal horn images at 20X in WT and *Fire*$^{\Delta/\Delta}$ mice, scale bar = 100 µm.
(**B** and **C**) Perivascular macrophages (IBA1$^{+}$ CD45$^{hi}$) in the spinal dorsal horn are decreased in *Fire*$^{\Delta/\Delta}$ mice, while IBA1$^{-}$CD45$^{hi}$ cell densities remain unchanged (C) ($n_M$ = 3-4, $n_F$ = 5/group).
(**D** – **F**) Hotplate latencies to first nocifensive behaviour (D), hotplate nocifensive behaviours (E), and tail withdrawal latencies (F) following a 10 mg/kg acute morphine challenge in WT and *Fire*$^{\Delta/\Delta}$ mice treated chronically with morphine, separated by sex (n = 3-5/group).
Data are mean ± SEM; independent t test with Welch correction (B, C); two-way ANOVA with Holm-Sidak post-hoc (F); three-way repeated measures ANOVA (D, E). *$p<0.05$, **$p<0.01$, ****$p<0.0001$.
